# Pleiotropic role of PAX cyclolipopeptides in the *Xenorhabdus* bacterium mutualistically associated with entomopathogenic nematodes

**DOI:** 10.1101/2025.04.11.648461

**Authors:** Noémie Claveyroles, Anne Lanois-Nouri, Imane El Fannassi, Jean-Claude Ogier, Sylvie Pagès, Adrien Chouchou, Guillaume Cazals, Gilles Valette, Alyssa Carré-Mlouka, Alain Givaudan

**Author notes:** Correspondence and reprints. (N. Claveyroles), (A. Lanois), (I. El Fannassi) (J. C. Ogier), (S. Pagès), (A. Chouchou), (G. Cazals), (G. Valette), (A. Carré-Mlouka)* Correspondence and reprints, (A. Givaudan).

## Abstract

*Xenorhabdus* is an entomopathogenic bacterium involved in a mutualistic relationship with *Steinernema* nematodes. *Xenorhabdus* produces a multitude of specialized metabolites by Non-Ribosomal Peptide Synthetases (NRPSs) pathways to mediate bacterial–nematode– insect interactions. PAX cyclolipopeptides are a family of NRP-type molecules whose ecological role remains poorly understood. In this study, the pleiotropic role of PAX peptides in the life cycle of *Xenorhabdus nematophila* has been investigated. By mass spectrometry analysis, we first demonstrated that PAX peptides were detected from the pathogenic stage up to the necrotrophic stage. We discovered that the bromothymol blue adsorption phenotype historically used to discriminate *Xenorhabdus* variants was associated with the presence of PAX peptides. We found that PAX peptides were positively involved in biofilm formation and negatively involved in swimming motility. PAX peptides were also shown to promote *in vivo* the production of infective *Steinernema* juveniles, suggesting their involvement in the mutualistic relationship between *Xenorhabdus* and its nematode partner. Finally, we showed that the *paxTABC* cluster as well as PAX peptide production was conserved across the whole *Xenorhabdus* genus except in *Xenorhabdus poinarii* and *Xenorhabdus ishibashii*. This work has revealed multiple new ecological roles for NRP-type peptides.

**Importance:** *Xenorhabdus* bacteria are models of particular interest for their mutualistic relationship with *Steinernema* nematodes and their ability to produce a wide range of natural NRP-type bioactive metabolites. These compounds are mostly studied for their medical or industrial applications, but their ecological role is poorly understood. This study provides a dynamic characterization of PAX cyclolipopeptide presence during *Xenorhabdus nematophila* life cycle, as well as confirmation of their production by 7 different strains within the *Xenorhabdus* genus. We revealed new multiple functions for PAX peptides in biofilm formation, swimming motility and juvenile nematode production. A deeper understanding of how PAX peptides interact with the nematode host would provide a better insight into the role of these cyclolipopeptides in bacterial-nematode mutualism.

## Introduction

Entomopathogenic bacteria of the genus *Xenorhabdus* (*Morganellaceae*) are hosted by nematodes of the *Steinernema* genus, with which they have co-evolved to establish long-lasting, mutually beneficial interactions (Goodrich-Blair *et al*., 2007). *Xenorhabdus* are carried into the insect larvae by the infective juvenile (IJs) stage of nematodes. IJs are living in the soils seeking insect larvae to infest. Once they have found a prey, IJs penetrate insect larvae through their natural orifices and release their *Xenorhabdus* bacteria into the insect hemocoel (Sicard *et al*., 2004). Bacteria cause septicemia and death of the insect larvae within 24 to 48 hours, enabling the nematodes to use the insect cadaver as a nutritive resource and a host for their development (Stock *et al*., 2019). Several nematode reproduction cycles occur successively inside the cadaver (Nguyen *et al*., 2007). During this necrotrophic stage, *Xenorhabdus* and *Steinernema* nematodes interact with a range of microorganisms from the nematode microbiota (Frequently Associated Microbiota, FAM) and the insect microbiota (Walsh *et al*., 2003 ; Singh *et al*., 2014 ; Ogier *et al*., 2020). When nutrients are depleted, mutualistic bacteria and nematodes reassociate and IJs emerge in the soil seeking a new host. Entomopathogenic nematodes are considered as promising biological control agents of key insect pests (Lacey *et al*., 2015).

To cope with changing conditions during its complex life cycle, *Xenorhabdus* can produce a wide variety of specialized metabolites including Non-Ribosomal Peptides (NRPs) (Tobias *et al*., 2017; Shi *et al*., 2022). Non-Ribosomal Peptide Synthetases (NRPSs) are modular enzymes that catalyze synthesis of a wide range of peptides from a variety of proteinogenic and non-proteinogenic amino acid substrates. In *Xenorhabdus nematophila*, the *ngrA* gene encodes the phosphopantetheinyl transferase (PPTase), an enzyme that activates NRPS by transferring a cofactor (4’-phosphopantetheine), thus enabling the production of NRP-type metabolites (Beld *et al*., 2014). It has been shown that *ngrA*-dependant compounds are responsible for most of the antimicrobial activity of *X. nematophila* and are required to eliminate bacterial competitors *in vitro* (Singh *et al*., 2015). It has also been demonstrated that *ngrA*-dependent compounds from *X. nematophila* are required for optimal growth and development of their nematode partner *in vivo*. Indeed, *Steinernema carpocapsae* reproduction was reduced in insects infected with a *ngrA* mutant (Singh *et al*., 2015). Similar results have been observed with *Photorhabdus luminescens* associated with the nematode *Heterorhabditis bacteriophora* (Ciche *et al*., 2001).

Studies on NRP metabolites in *Xenorhabdus* have so far focused mainly on their antimicrobial potential (Booysen *et al*., 2020). These include xenocoumacins, odilorhabdins, xenoamicins, xenematides, xenortides, rhabdopeptides, RXPs, GameXpeptides, nematophin, xenobactin, szentiamide, bicornitun, taxlllaids A-G and PAX peptides (McInerney *et al*., 1991 ; Pantel *et al*., 2018 ; Zhou *et al*., 2013 ; Lang *et al*., 2008 ; Reimer *et al*., 2014 ; Hacker *et al*., 2018 ; Cai *et al*., 2017 ; Bode *et al*., 2012 ; Li *et al*., 1997 ; Grundmann *et al*., 2013 ; Ohlendorf *et al*., 2011 ; Fuchs *et al*., 2012 ; Kronenwerth *et al*., 2014 ; Gualtieri *et al*., 2009). Some of these compounds also exhibit insecticidal (xenocoumacins (Touray *et al*., 2024) ; xenematides (Lang *et al*., 2008) ; RXPs (Cai *et al*., 2023)), antiprotozoal (xenoamicins (Zhou *et al*., 2013); xenortides (Reimer *et al*., 2014) ; RXPs (Cai *et al*., 2017) ; xenobactin (Grundmann et al., 2013) ; taxlllaids A-G (Kronenwerth *et al*., 2014)), acaricide (xenocoumacins (Incedayi *et al*., 2021)), and anti-inflammatory (xenocoumacins (Erkoc *et al*., 2021)) properties.

Our study is focusing on the cyclolipopeptides PAX (Peptide Antimicrobial from *Xenorhabdus*) family (Gualtieri *et al*., 2009). NRPS enzymes involved in PAX peptide biosynthesis are encoded by the *paxTABC* gene cluster. The *paxA*, *paxB* and *paxC* genes encode the three NRPS enzymes responsible for heptapeptide synthesis and cyclization. The *paxT* gene encodes a putative membrane transport protein suspected of exporting PAX peptides outside the cell. PAX cyclolipopeptides are cationic peptides containing 7 amino acids, among which 5 lysines constitute a cycle, with a N-terminal fatty acid moiety (Fuchs *et al*., 2011). Different PAX peptides, with variations in the length of the fatty acid chain or the nature of the amino acid in position 2 (lysine or arginine), have been identified among different strains and species of *Xenorhabdus* (Gualtieri *et al*., 2009; Fuchs *et al*., 2011; Dreyer *et al*., 2019). Bacterial specialized metabolites are widely investigated in the literature for their medical or agricultural applications, but their ecological roles are still poorly understood (Schmidt *et al*., 2019). PAX peptides were first studied for their antimicrobial activities against *Microccocus luteus* and *Fusarium oxysporum* (Gualtieri *et al*., 2009). PAX peptides exhibit no insecticidal activity against *Spodoptera littoralis*, *Galleria mellonella*, *Locusta migratoria*, *Manduca sexta* and insect hemocytes, nor nematicidal activity against *Caenorhabditis elegans* (Gualtieri *et al*., 2009; Abebew *et al*., 2022). As external addition of synthetic labelled PAX peptides to *Xenorhabdus doucetiae* resulted in their localization at the cell surface, it was suggested that PAX peptides may provide protection against cationic antimicrobial peptides (AMPs) produced by the insect via positive charge repulsion mechanisms (Vo *et al*., 2021).

In this study, we revealed by mass spectrometry that PAX peptides were detected from the pathogenic stage to the late necrotrophic stage of *X. nematophila* life cycle. We therefore aimed to elucidate the role(s) of PAX peptides using two deletion mutants of *X. nematophila* F1, Δ*paxA* and Δ*ngrA* mutants (Lanois *et al*, 2022). By comparing the two mutants with the wild-type (WT) strain, we demonstrated different phenotypic characteristics on agar media, in swimming motility and in biofilm formation *in vitro*. By minimum inhibitory concentration (MIC) assays, we showed that PAXs were weakly antimicrobial towards natural competitors of *Xenorhabdus*. We also assessed the pathogenicity of bacteria against insect larvae. Using aposymbiotic nematodes reassociated with the WT strain or the Δ*paxA* or Δ*ngrA* mutants, we investigated the involvement of PAX peptides in nematode reproductive success *in vivo*. Finally, we have highlighted the distribution of *paxTABC* genes and detected PAX peptide production throughout the *Xenorhabdus* genus. These findings reveal that PAX peptides are highly conserved compounds playing multiple roles in *Xenorhabdus* life cycle.

## Results

### PAX peptides are mostly detected during the necrotrophic stage of *X. nematophila* life cycle

To understand the importance of PAX peptides in the life cycle of *X. nematophila* F1, PAX peptide presence was investigated at each stage of the bacterial life cycle. We first characterized by MALDI-TOF-MS all PAX peptides produced on stationary-phase extracts of the WT strain in 48 h liquid culture. Twelve PAX peptides previously identified in the literature by Fuchs *et al*., 2011 were detected in stationary-phase-extracts of the WT strain: PAX1’ (m/z=1052.772 Da), PAX3’ (1066.787), PAX2’/PAX7 (1080.783), PAX4’ (1094.797), PAX6 (1078.766), PAX8 (1102.772), PAX9 (1106.797), PAX10 (1108.812), PAX11 (1120.808), PAX12 (1122.818), PAX13 (1134.828) (Table S1).

Then, the presence of most frequently detected PAX peptides (PAX1’, PAX2’/PAX7, PAX3’ and PAX6) was analyzed at 20 h, 48 h, 72 h, 5 days and 7 days under three different experimental setups: i) *in vitro* bacterial cultures, ii) insect larvae infected with *X. nematophila* WT bacteria, iii) insect larvae infested with *S. carpocapsae* SK27 nematodes carrying *X. nematophila* WT bacteria (Figure 1, Table S2). An extraction control was performed by adding a synthetic PAX derivative (DP18, Figure S1) before the extraction steps. The m/z values (+/-0.1 Da) of selected PAX peptides and the extraction control DP18, were screened by MALDI-TOF-MS analysis. In bacterial culture in LB culture medium, PAX peptides were constantly found from 20 h (late exponential stage, Figure S2) up to 7 days. Under *in vivo* conditions of *Galleria mellonella* larvae infected with bacteria or infested with nematodes, higher variability was observed at 20 h between biological samples. However, PAX peptides were detected from 48 h to 7 days in both *in vivo* conditions, except for PAX6, which is less frequently found in bacterial-infected larvae. These results demonstrate that PAX peptides are detected from the end of the pathogenic phase (20 h) to the late necrotrophic phase (7 days) of the *Xenorhabdus* life cycle.

**Figure 1:**
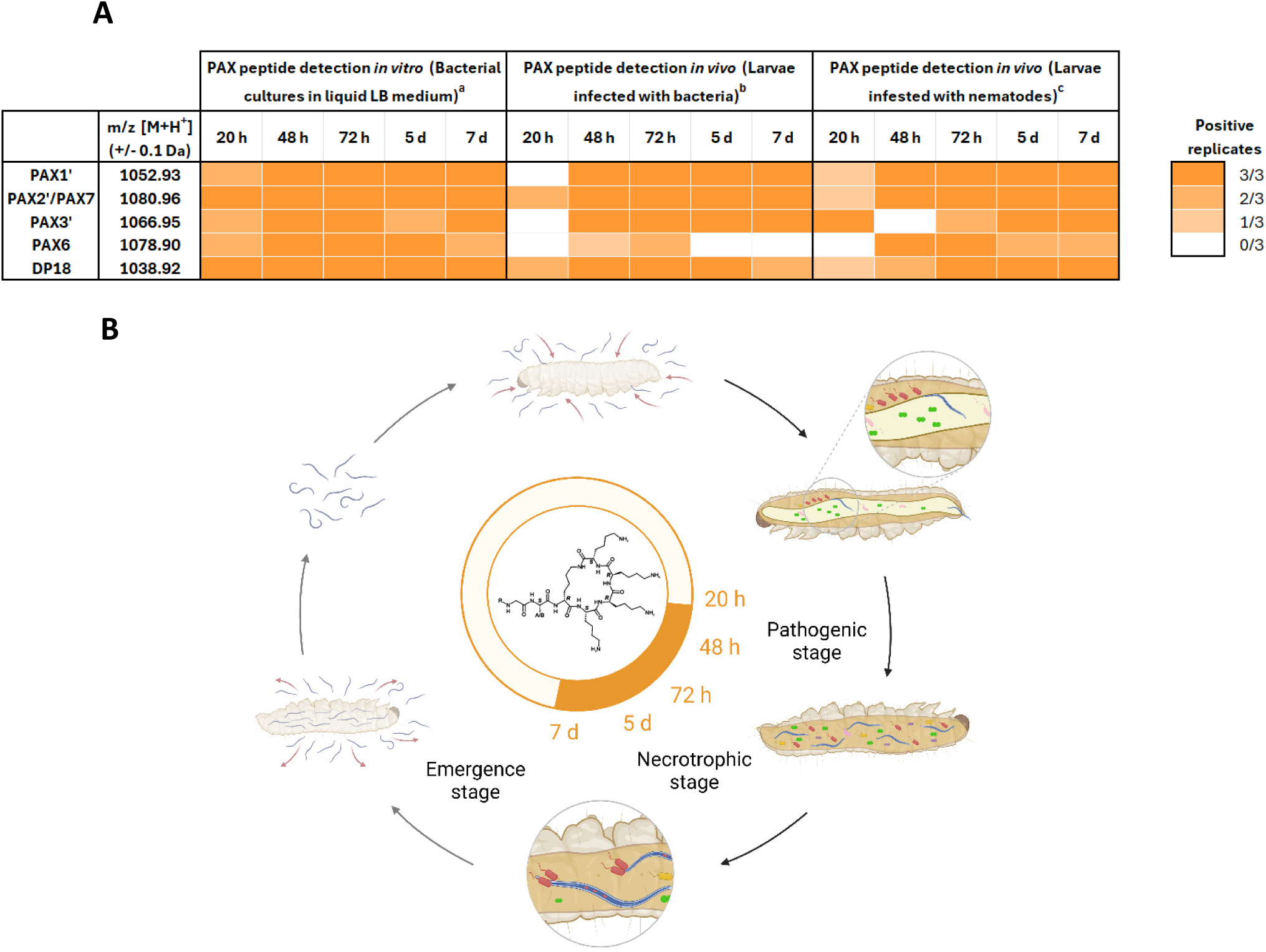
Presence of PAX peptides in extracts of *X. nematophila* WT grown *in vitro* and *in vivo*. A) Detection by MALDI-TOF-MS analysis of selected PAX peptides (PAX1’, PAX2’/PAX7, PAX3’, PAX6) and the extraction control DP18, in extracts of *X. nematophila* WT grown *in vitro* in liquid LB medium (^a^) and *in vivo* in 10 *G. mellonella* larvae injected with *X. nematophila* WT (^b^) or infested with *Steinernema carpocapsae* SK27 IUs nematodes (^c^). A synthetic PAX peptide derivative DP18 (m/2=1038.92 +/-0.1 Da) was added as a positive extraction control before the extraction steps. Experiments were performed in triplicate. Orange squares indicate the number of positive biological replicates in which the m/2 value (+/-0.1 Da) of the corresponding PAX peptides was detected. The corresponding mass spectra are given in Table S2. B) Visual representation of PAX peptide presence in different stages of *Xenorhabdus* life cycle. The orange circle area represents the period of detection of PAX peptides. Figure created with Biorender.

### *paxA* mutation affects lecithinase-like activity, lipolytic activity and bromothymol blue adsorption

To assess the impact of PAX peptide production on phenotypic characteristics, a deletion mutant (Δ*paxA*) was constructed in *X. nematophila* F1 by inserting an Ω-cam interposon by allelic exchange at the beginning of the *paxA* gene, the first NRPS gene in the *paxTABC* cluster. As phosphopantetheinyl transferase (PPTase) enzymes enable NRPS megasynthetase activation, a PPTase deficient mutant (Δ*ngrA*) of *X. nematophila* F1 was also used as a global non-producer of NRP-type metabolites. The absence of PAX peptide production by Δ*paxA* (Figure 2A) and Δ*ngrA* (Figure 2B) mutants was confirmed by MALDI-TOF-MS in extracts from stationary-phase cultures.

**Figure 2:**
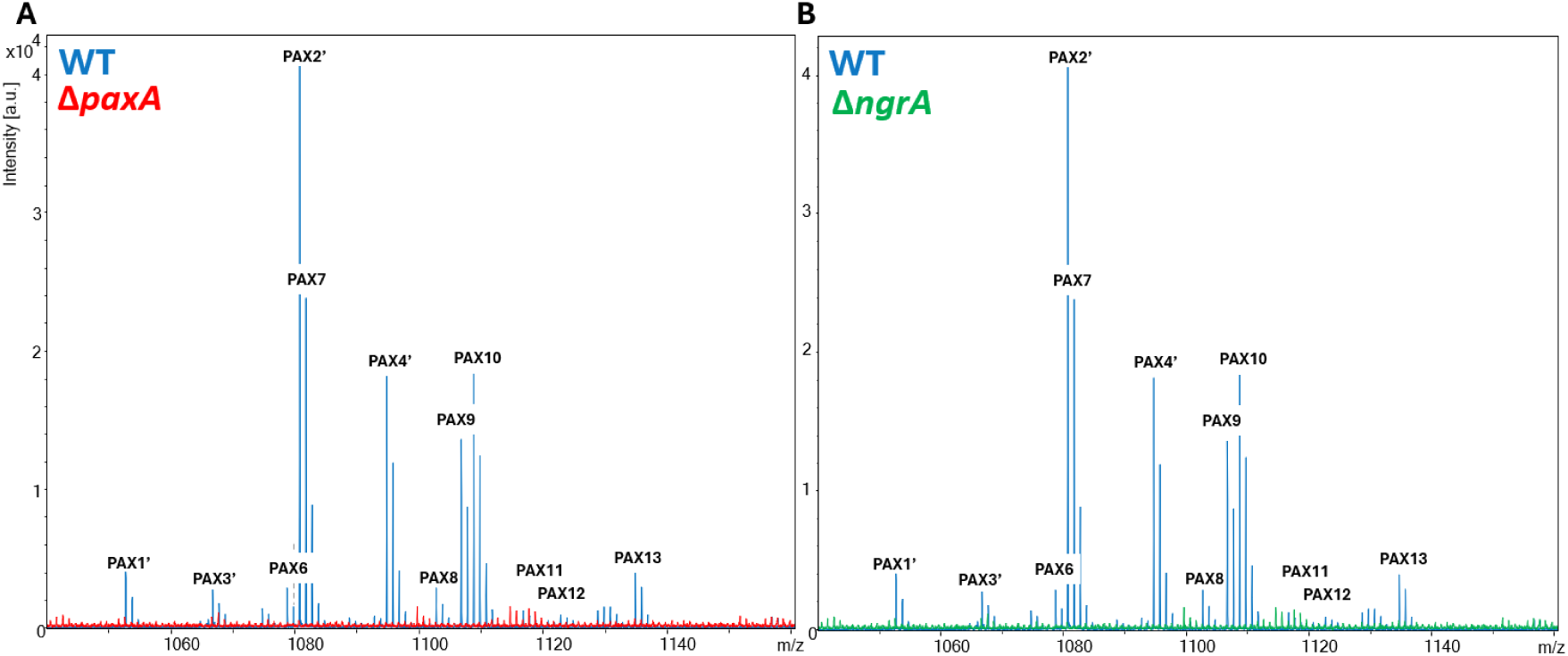
Presence of PAX peptides in extracts of *X. nematophila* WT (blue), Δ*paxA* (red) and Δ*ngrA* (green). **(A)** Merging of MALDI-TOF-MS mass spectra of extracts from 48 h stationary-phase cultures of WT and Δ*paxA* strains. **(B)** Merging of MALDI-TOF mass spectra of extracts from 48 h stationary-phase cultures of WT and Δ*ngrA* strains. The monoisotopic m/2 values of PAX peptides are shown with their numbers previously identified by Fusch *et al*., 2011. Peak heights represent intensity in arbitrary units (a.u). Exact m/2 values (+/-0.05 Da) of each PAX peptide are listed in Table S1.

The phenotypic traits of the Δ*paxA* and Δ*ngrA* mutants were then tested through various phenotypic assays on agar media. The Δ*paxA* and Δ*ngrA* mutants displayed enhanced lipolytic activity on Tween 20, 40, 60, 80 and 85 and suppressed lecithinase-like activity compared to the WT strain (Table 1). However, no difference in haemolysis was observed. As expected, the Δ*ngrA* mutant showed weaker antimicrobial activity against *M. luteus* than the WT strain or Δ*paxA* mutant. Surprisingly, both mutant strains displayed red colonies on NBTA medium, indicating no bromothymol blue (BBT) adsorption (Figure S3). These observations demonstrate that PAX peptides are responsible for BBT adsorption. Together, these results indicate that the mutation of Δ*paxA* and Δ*ngrA* induce pleiotropic effects on phenotypes of *X. nematophila* similar for both mutants, excepting concerning antimicrobial activity against *M. luteus*.

**Table 1:**
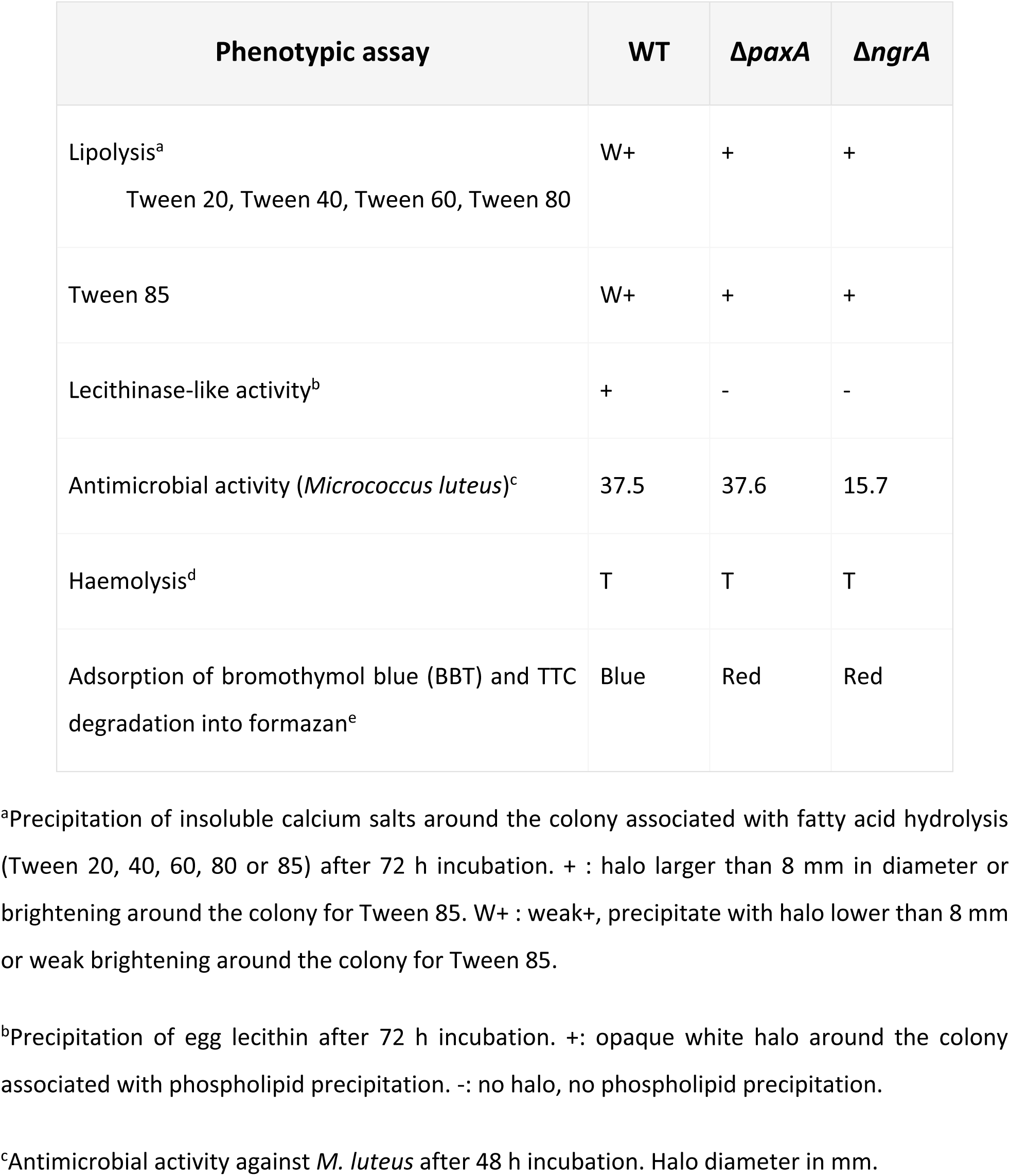
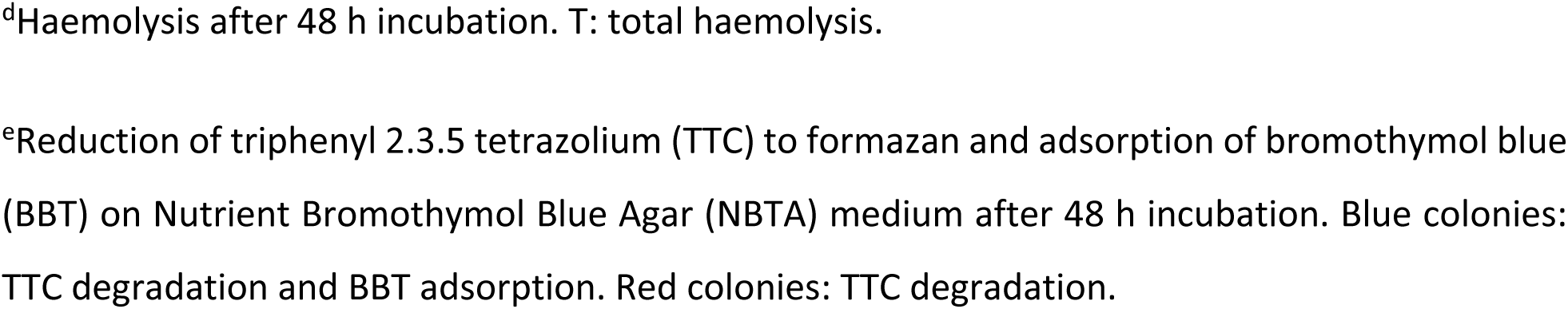
Phenotypic traits of *X. nematophila* WT, Δ*paxA* and Δ*ngrA* on agar media.

| Phenotypic assay | WT | $\Delta paxA$ | $\Delta ngrA$ |
| --- | --- | --- | --- |
| Lipolysis<sup>a</sup><br>Tween 20, Tween 40, Tween 60, Tween 80 | W+ | + | + |
| Tween 85 | W+ | + | + |
| Lecithinase-like activity<sup>b</sup> | + | - | - |
| Antimicrobial activity (*Micrococcus luteus*)<sup>c</sup> | 37.5 | 37.6 | 15.7 |
| Haemolysis<sup>d</sup> | T | T | T |
| Adsorption of bromothymol blue (BBT) and TTC degradation into formazan<sup>e</sup> | Blue | Red | Red |

<sup>a</sup>Precipitation of insoluble calcium salts around the colony associated with fatty acid hydrolysis (Tween 20, 40, 60, 80 or 85) after 72 h incubation. + : halo larger than 8 mm in diameter or brightening around the colony for Tween 85. W+ : weak+, precipitate with halo lower than 8 mm or weak brightening around the colony for Tween 85.
<sup>b</sup>Precipitation of egg lecithin after 72 h incubation. +: opaque white halo around the colony associated with phospholipid precipitation. -: no halo, no phospholipid precipitation.
<sup>c</sup>Antimicrobial activity against *M. luteus* after 48 h incubation. Halo diameter in mm.
<sup>d</sup>Haemolysis after 48 h incubation. T: total haemolysis.
<sup>e</sup>Reduction of triphenyl 2.3.5 tetrazolium (TTC) to formazan and adsorption of bromothymol blue (BBT) on Nutrient Bromothymol Blue Agar (NBTA) medium after 48 h incubation. Blue colonies: TTC degradation and BBT adsorption. Red colonies: TTC degradation.

### PAX peptides are involved in biofilm formation and swimming motility

Biofilm formation, swimming motility and swarming motility were investigated *in vitro* for *X. nematophila* WT, Δ*paxA* and Δ*ngrA* strains (Figure 3). The Δ*paxA* (*p*<0.05) and Δ*ngrA* (*p*<0.01) mutants showed significantly lower biofilm formation than the WT strain (Figure 3A). Conversely, Δ*paxA* (*p*<0.001) and Δ*ngrA* (*p*<0.01) mutants showed significantly enhanced swimming motility on LB agar 0.35% medium compared with the WT strain (Figure 3B). No difference in swarming motility was observed between the three strains onto 0.7% Eiken agar plate (Figure 3C). PAX peptides are therefore positively involved in biofilm formation and negatively in swimming motility.

**Figure 3:**
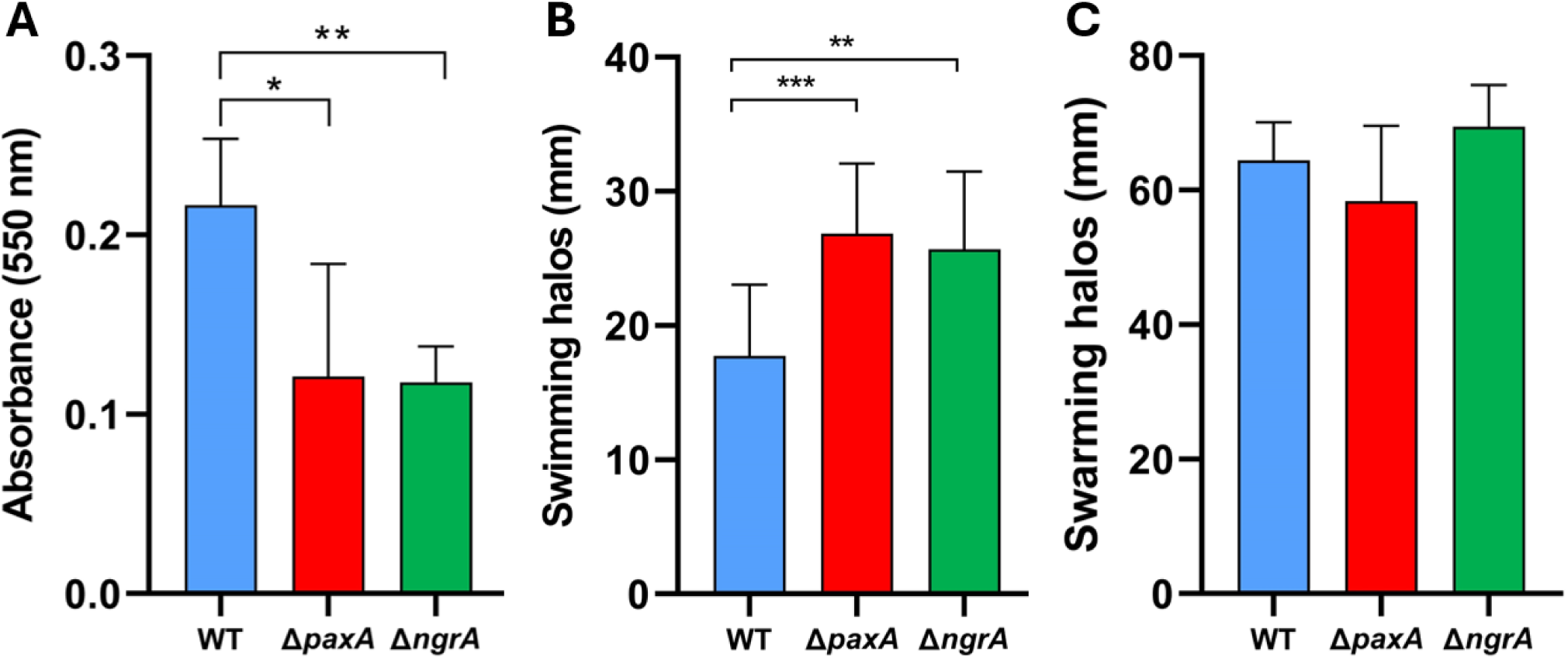
Biofilm formation, swimming and swarming motilities of *X. nematophila* WT (blue), Δ*paxA* (red) and Δ*ngrA* (green). **A)** Biofilm formation was assessed by 1% crystal violet staining of 48 h static bacterial cultures and quantification of absorbance at 550 nm. Student t test, n=4. Swimming motility **(B)** and swarming **(C)** were assessed after dropping 5 μL of overnight bacterial culture onto LB 0.35% agar (Student’s t test, n=4) and LB 0.7% agar Eiken (Wilcoxon-Mann-Whitney test, n=4) media, respectively. Halo diameter was measured after 18h of growth. *p<0.05, \*\**p*<0.01, \*\*\**p*<0.001.

### PAX peptides are slightly involved in bacterial virulence towards insects

To determine whether PAX peptides were involved in the virulence of *X. nematophila in vivo*, pathogenicity assays of *X. nematophila* WT, Δ*paxA* and Δ*ngrA* strains were carried out by injection into *S. littoralis* larvae (Figure 4). A weak difference (1h40, *p*<0,05) of larval mortality between the Δ*paxA* mutant and the WT strain was observed (Figure 4A), but no difference between the Δ*ngrA* mutant and the WT strain (Figure 4B). In a control experiment, we showed that the WT and mutant strains displayed no growth difference when cultivated in LB (Figure S2).

**Figure 4:**
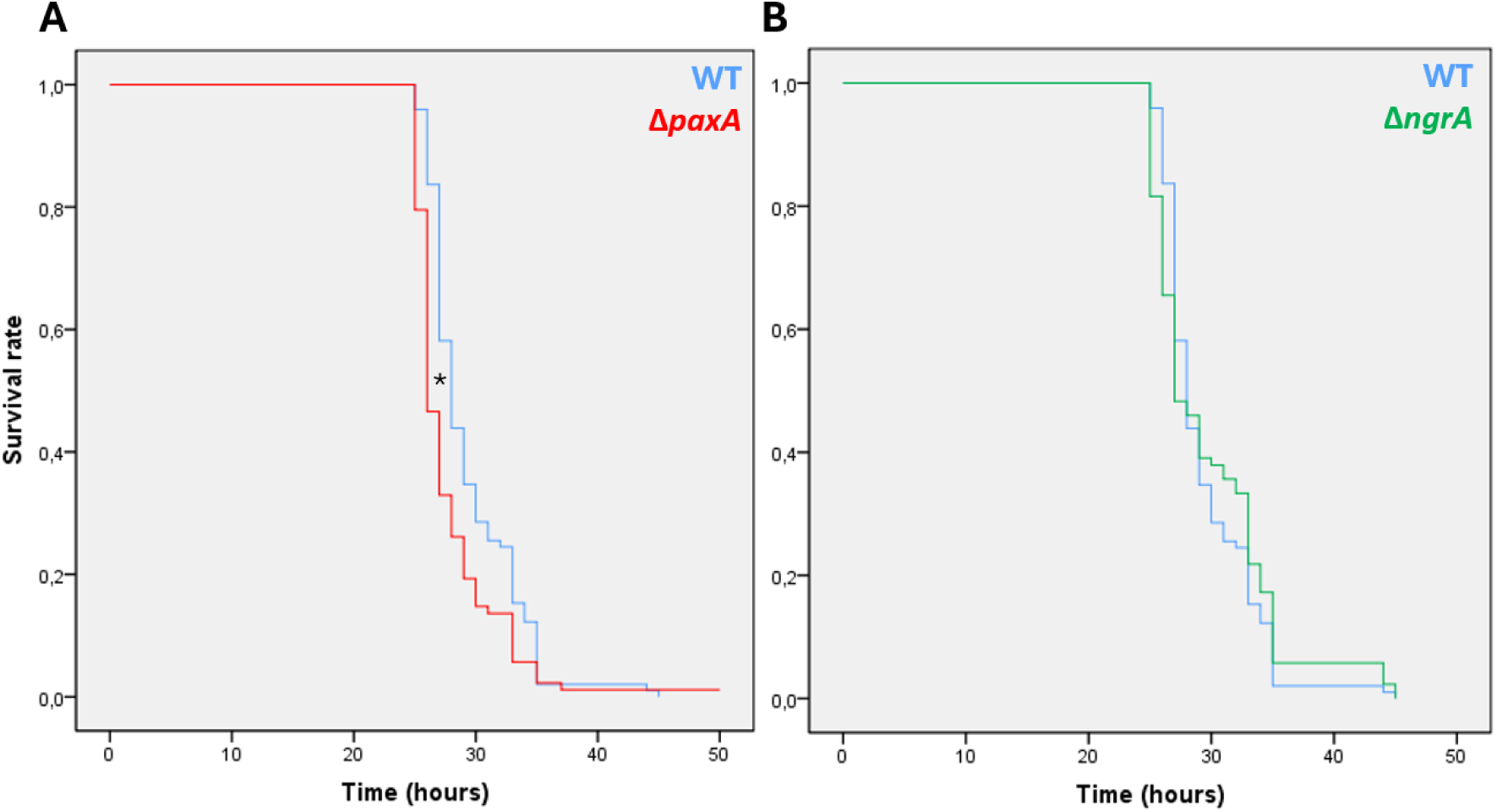
Pathogenicity assay of *X. nematophila* WT (blue), Δ*paxA* (A, red) and Δ*ngrA* (B, green) towards *S. littoralis* larvae (L6 stage). Kaplan-Meier survival curves were generated with Statistical Package for the Social Sciences V18.0 (SPSS). Each insect was injected with 1000–3000 CFUs of the respective strains. Wilcoxon-Mann-Whitney tests, n=80, *p<0.05.

To test the hypothesis of a role for PAX peptides in the resistance of *X. nematophila* to insect AMPs (Vo *et al*., 2021), MIC assays were carried out using various insect AMPs (cecropin A, cecropin B1, cecropin C, cecropin P1, drosocin) against the WT, Δ*paxA* and Δ*ngrA* strains. MIC assays were also carried out using bacterial AMPs (colistin, polymyxin B, NOSO 95-C, synthetic PAX1’, synthetic PAX2’ and synthetic PAX7) to determine whether PAX peptides would also enable resistance to competitive bacteria within the insect cadaver (Table 2, Figure S1). No clear difference in MIC was detected between the WT and mutant strains, for any of the bacterial or insect AMPs tested. In our experiments, PAX peptides are not involved in resistance of *X. nematophila* to the tested insect and bacterial AMPs.

**Table 2:** Minimum inhibitory concentration (MIC in µg. mL^-1^) assays of insect and bacterial antimicrobial peptides on *X. nematophila* WT, Δ*paxA* and Δ*ngrA*.

| Antimicrobial peptide | WT | $\Delta\text{paxA}$ | $\Delta\text{ngrA}$ |
| --- | --- | --- | --- |
| Cecropin A | 12.5 | 12.5 | 10.5 |
| Cecropin B1 | 6.25 | 6.25 | 6.25 |
| Cecropin C | 12.5 | 12.5 | 25 |
| Cecropin P1 | >12.5 | >12.5 | 12.5 |
| Drosocin | >50 | >50 | >50 |
| Colistin | 3.9 | 1.95 | 3.9 |
| Polymyxin B | 0.245 | 0.245 | 0.245 |
| NOSO 95-C (Odilorhabdin) | > 1700 | > 1700 | > 1700 |
| Synthetic PAX1' | >1500 | >1500 | >1500 |
| Synthetic PAX2' | 187.5 | 375 | 375 |
| Synthetic PAX7 | 187.5 | 187.5 | 375 |

### Microorganisms inhabiting the ecological niche of *Xenorhabdus* exhibit low susceptibility to PAX peptides

To determine whether the production of PAX peptides could enable *Xenorhabdus* to outcompete the microorganisms present in the insect cadaver, MIC assays were performed using synthetic PAX1’, PAX2’ and PAX7 (Table 3, Figure S1). For this purpose, 10 microorganisms from the microbiota (FAM) of *S. carpocapsae* nematodes (*Stenotrophomonas maltophilia* ALL5, *Pseudomonas protegens* PPSC10, *Ochrobactrum* sp. ALL4, *Achromobacter* sp. D7.1, *Alcaligenes faecalis* SC, *Pseudochrobactrum* sp. AL3, *Brevundimonas* sp. ALL3) and from the microbiota of the insect *Spodoptera* (*Enterococcus mundtii* SP, *Diutina rugosa* GC) were selected. *M. luteus* CIP103430 was also used as a positive control for the antimicrobial activity of synthetic PAX peptides, as it had already been identified as susceptible to PAX peptides (Gualtieri *et al*., 2009). Microorganisms from the nematode microbiota appear to exhibit low susceptibility to synthetic PAX tested with a minimum MIC of 31.25 µg. mL^-1^ for PAX1’ and PAX7 against *Brevundimonas* sp.. *E. mundtii* and *D. rugosa* (insect microbiota) show higher susceptibilities, with MICs down to 15.63 µg. mL^-1^ for PAX7 against *E. mundtii* and 25 µg. mL^-1^ for PAX1’ against *D. rugosa*. The antimicrobial effect of synthetic PAX peptides against the microorganisms co-occurring with *Xenorhabdus* in the insect cadaver niche is moderate.

**Table 3:** Minimum inhibitory concentration (MIC in µg. mL^-1^) assays of synthetic PAX1’, PAX2’ and PAX7 on selected microorganisms from nematode microbiota (*Steinernema carpocapsae*, pink) and insect microbiota (*Spodoptera* larvae, yellow).

| Microorganisms | Synthetic PAX1' | Synthetic PAX2' | Synthetic PAX7 |
| --- | --- | --- | --- |
| *Micrococcus luteus* CIP103430 | 3.9 | 1.95 | 0.49 |
| *Stenotrophomonas maltophilia* ALL5 | >250 | 125 | 125 |
| *Pseudomonas protegens* PPSC10 | >250 | 125 | 62.5 |
| *Ochrobactrum* sp. ALL4 | >250 | 62.5 | 62.5 |
| *Achromobacter* sp. D7.1 | >250 | 62.5 | 62.5 |
| *Alcaligenes faecalis* SC | >250 | 125 | 125 |
| *Pseudochrobactrum* sp. AL3 | >250 | 125 | 62.5 |
| *Brevundimonas* sp. ALL3 | 31.25 | 62.5 | 31.25 |
| *Enterococcus mundtii* SP | 125 | 31.25 | 15.63 |
| *Diutina rugosa* GC | 25 | 62.5 | 62.5 |

### PAX peptides enhance of infective juvenile nematode production

Since PAX peptides were detected during the necrotrophic stage (Figure 1), we questioned whether PAX are involved in the production of IJ nematodes. *G. mellonella* larvae were infested with *S. carpocapsae* SK27 reassociated with the WT strain, Δ*paxA* or Δ*ngrA* mutants (Figure 5). A difference in reproductive success was observed between the WT strain and both mutant strains (n=14 to 19) at the end of the 1st generation (Figure 5A), then between the WT strain and the Δ*paxA* mutant (n=60) at the end of the 2nd generation (Figure 5B, Wilcoxon-Mann-Whitney tests, *p*<0.05), and delayed emergence were noticed in both mutants (Figure S4). The second generation selects the best-performing IJs and reduces the differences observed between strains. PAX peptides are therefore involved in the production of *S. carpocapsae* IJs *in vivo*.

**Figure 5:**
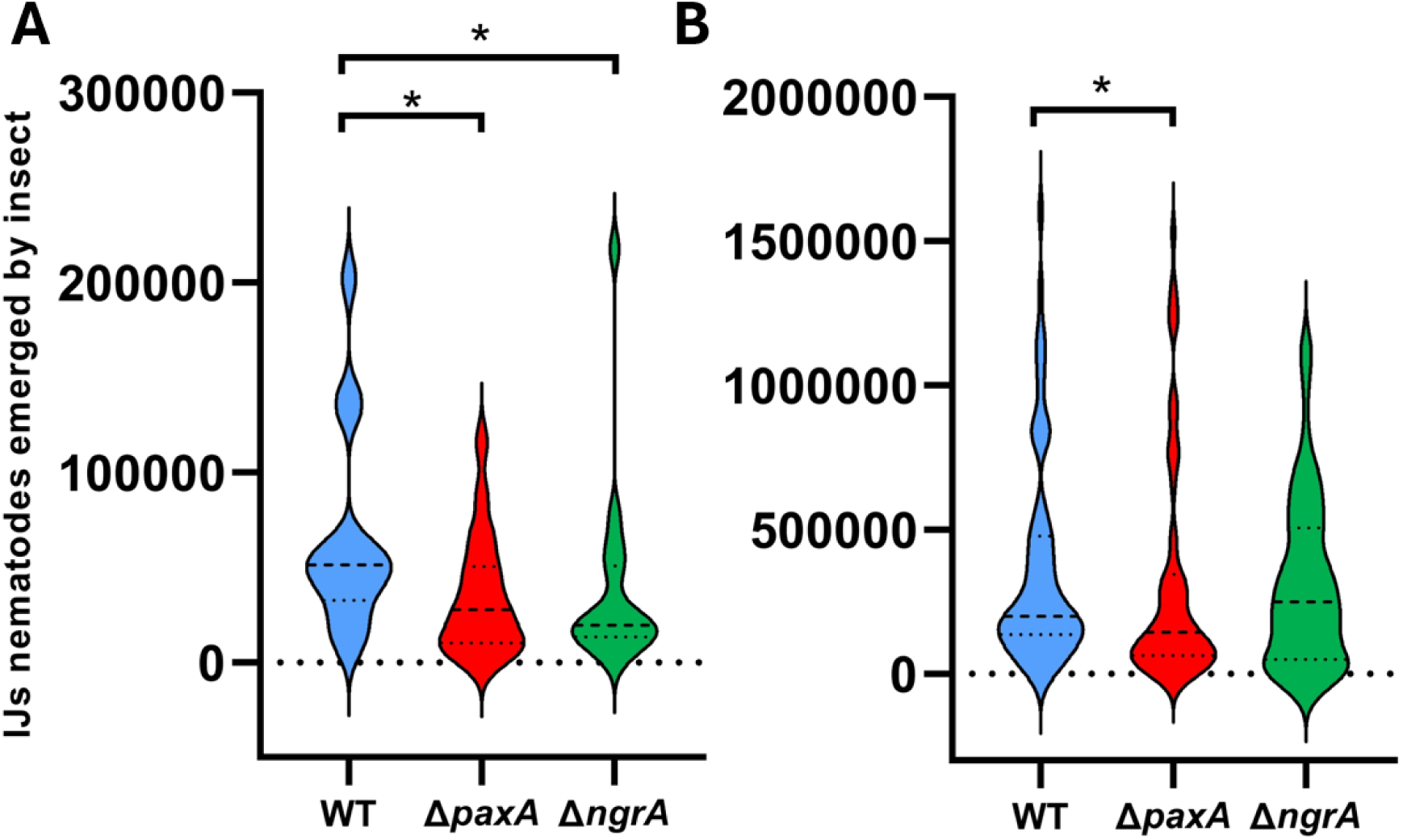
Reproductive success of aposymbiotic *Steinernema carpocapsae* SK27 nematodes reassociated with *X. nematophila* WT, Δ*paxA* or Δ*ngrA* in *G. mellonella* larvae. Emerging Infective Uuveniles (IUs) nematodes were counted at 15- and 30-days post-mortem and cumulated. The total number of IUs emerging per insect is represented at the end of the 1st generation (n=14 to 19) **(A)** and 2nd generation (n=60), on three independent experiments **(B)**. Wilcoxon-Mann-Whitney test, *p<0.05.

### PAX peptide-producing ability is widespread within the *Xenorhabdus* genus

To determine whether PAX peptide production ability is conserved within the *Xenorhabdus* genus, we first studied the distribution and the genetic environment of the *paxTABC* genes in 40 strains of *Xenorhabdus* belonging to 23 species compared to *X. nematophila* F1 (Figure 6). The surrounding genes and their organization are conserved among *X. nematophila* species, especially the *nilABC* genes, but not in other *Xenorhabdus* species. For NRPS genes, modules (*paxA* (m1), *paxB* (m2, m3, m4), *paxC* (m5, m6, m7)) were analysed independently as functional entities. *paxTABC* genes are present in almost all studied *Xenorhabdus* strains excepting *X. poinarii* G6 and *X. ishibashii* DSM 22670. Although the entire cluster is well distributed among the *Xenorhabdus* strains, the *paxC* gene (m5, m6 and m7) is absent in a few genomes (*Xenorhabdus szentirmaii* DSM 16338, *Xenorhabdus kozodoii* DSM 17907 and *Xenorhabdus beddingii* DSM 4764). Pairwise alignements showed that the *paxTABC* cluster exhibits high conservation across all strains within *Xenorhabdus bovienii* species and within *X. nematophila* species but shows lower identity between the two species (Figure S5). Thus, the *paxTABC* cluster is present throughout the entire *Xenorhabdus* genus, but with variability among NRPS modules.

**Figure 6:**
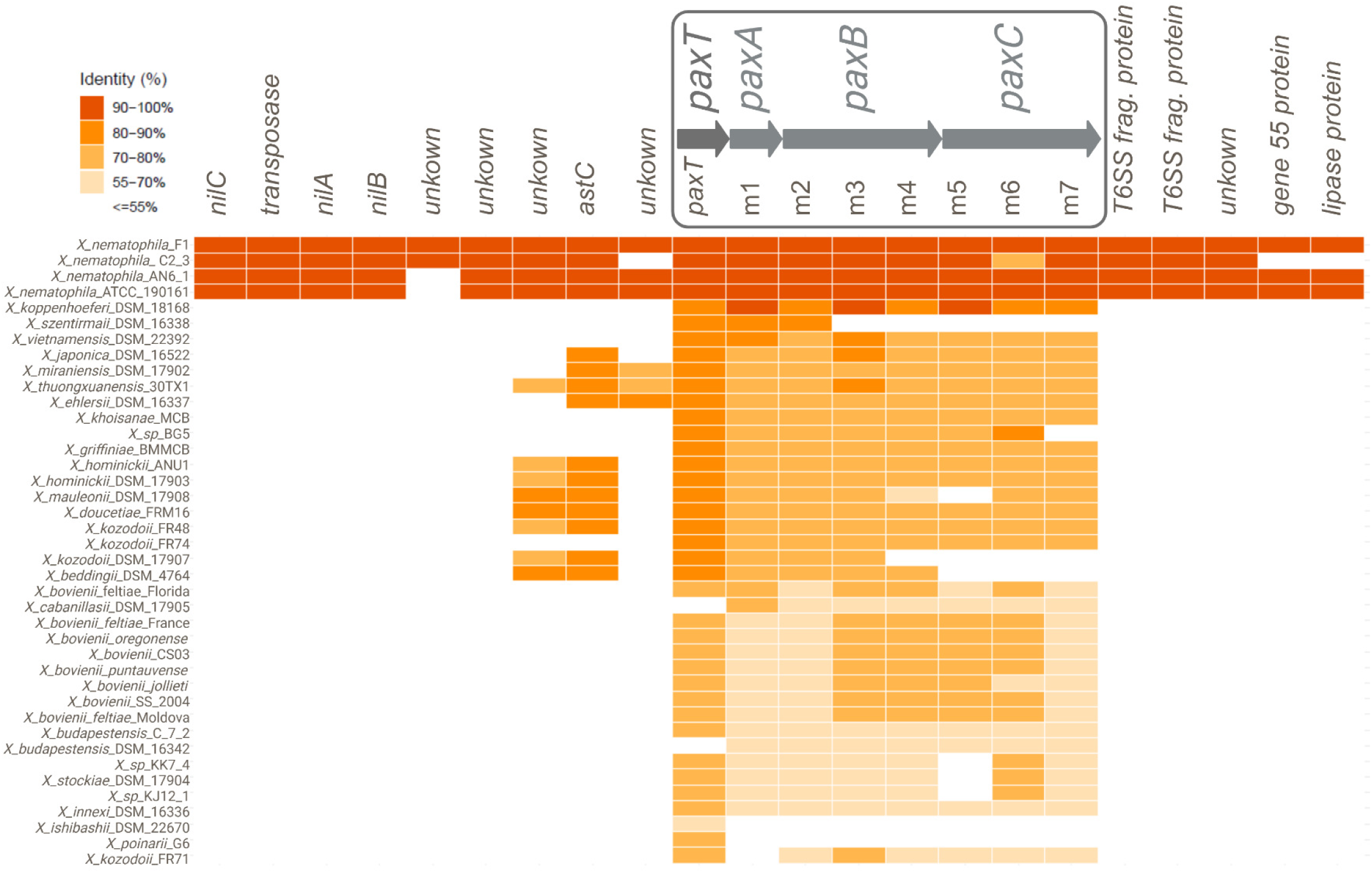
Distribution and genetic environment of *paxTABC* among the *Xenorhabdus* genus. The *paxTABC* and surrounding genes of *X. nematophila* F1 were aligned with those of 40 strains belonging to 23 *Xenorhabdus* species by NCBI BLAST+ blastn. The NRPS genes within this cluster were specifically analy2ed by comparing their module nucleotide sequences (*paxA* : m1 ; *paxB* : m2, m3, m4 ; *paxC* : m5, m6, m7). Orange gradient squares represent the nucleotide percentage identity of genes and modules for each orthologous sequence compared to their counterpart of *X. nematophila* F1. The *paxTABC* operon is shown at the top of the figure, along with its surrounding genes in *X. nematophila* F1 genome.

To correlate the presence of the *paxABC* genes with the actual production of PAX peptides, mass spectrometry analysis (MALDI-TOF-MS) were then performed on extracts from 48 h-cultures of 9 different strains representative of the main species of *Xenorhabdus* (Table 4, Table S3). The analysis focused on the m/z values (+/-0.1 Da) of the most detectable PAX peptides, PAX1’, PAX2’/PAX7, PAX3’, PAX6 and DP18 as the positive extraction control. The DP18 control was detected in all samples. At least one of the PAX peptides was detected in all tested strains, excepting *X. poinarii* G6 and *X. ishibashii* DSM 22670 (Table S3) for which no *paxTABC* cluster was evidenced (Figure 6). The most conserved PAX peptide in all producing strains is PAX1’. These results revealed that the PAX peptide-producing ability is widespread in the *Xenorhabdus* genus.

**Table 4:**
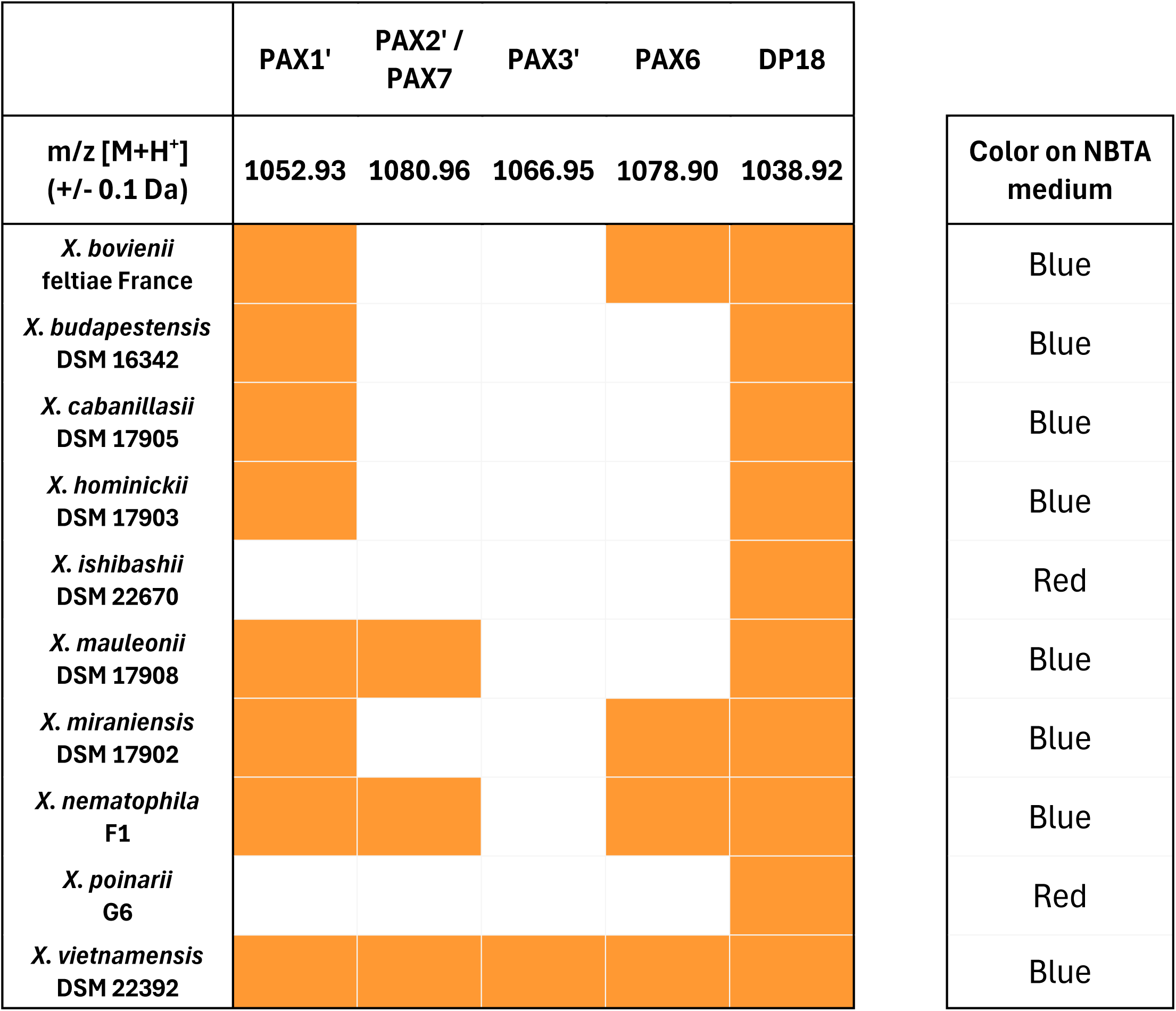
MALDI-TOF-MS detection of selected PAX peptide production in extracts from different strains of *Xenorhabdus* and colony color on NBTA medium. Orange squares indicate that the m/z value of the corresponding PAX peptide was detected at +/-0.1 Da. White squares mean that no m/z value corresponding to PAX peptides was detected. A synthetic PAX peptide derivative DP18 (m/z=1038.92 Da) was added as a positive extraction control before the extraction steps. The corresponding mass spectra are given in Table S3. Colony colors were assessed after streaking on NBTA medium and incubation for 48 h at 28°C.

To determine whether a correlation could be established between BBT adsorption and actual PAX peptide production, we examined the color of colonies grown on NBTA medium (Table 4). Similarly to the Δ*paxA* and Δ*ngrA* mutants (Figure S3), colonies of *X. poinarii* G6 and *X. ishibashii* DSM 22670, lacking the *paxTABC* cluster and for which no PAX peptides were detected in liquid culture, were red on NBTA medium. Therefore, the *Xenorhabdus* strains for which PAX peptide production was demonstrated were systematically observed as blue colonies on NBTA medium, whereas red colonies corresponded to strains unable to produce PAX peptides.

## Discussion

### BBT dye adsorption by bacterial colonies is a marker of PAX peptide production

PAX peptides constitute a family of positively charged cyclolipopeptides, presumably localized at the surface of *Xenorhabdus* bacteria (Vo *et al*., 2021). To investigate the role of PAX peptides, Δ*paxA* and Δ*ngrA* mutants were used. Mass spectrometry analysis confirmed the absence of PAX peptide production in both mutants, demonstrating that PAX peptide production in *Xenorhabdus* is *ngrA*-dependent. The NgrA PPTase is therefore likely involved in activation of the PaxABC synthetase, as well as other synthetases responsible for production of NRP metabolites and optimal antibiotic activities in *Xenorhabdus* (Beld *et al*., 2014 ; Singh *et al*., 2015 ; Ciezki *et al*., 2019).

Phenotypic characterization of laboratory cultures revealed similar defects in both Δ*paxA* and Δ*ngrA* mutant strains. This suggests that the changes observed in the NRPS-deficient Δ*ngrA* mutant, including the absence of lecithinase-like activity, increased Tween-lipase activity, abolition of BBT adsorption, reduced biofilm formation and increased swimming motility, are primarily due to the absence of PAX peptides. We showed that both mutants exhibited an enhanced precipitation zone in the agar containing Tween 20, 40, 60 or 80 and abolished lecithinase-like activity. Initially, lecithinase-like activity in *X. nematophila* F1 was partially characterized through the purification of five compounds that cause lecithin precipitates on agar, without exhibiting phospholipase C activity (Thaler *et al*., 1998). Given the absence of lecithin precipitates in the Δ*paxA* mutant, it is likely that these five compounds correspond to the five PAX peptides described by Gualtieri *et al*., 2009, who employed the same methodology.

Additionally, our study uncovered another striking difference between both mutants and the WT strain as colonies of both mutants displayed no adsorption of bromothymol blue dye on NBTA medium, unlike the WT strain. This phenotype has historically been used to distinguish phenotypic variants in *Xenorhabdus*, as WT strain shows blue colonies on NBTA medium, while secondary variants show red colonies (Akhurst, 1986). In addition, *X. nematophila* secondary variants are characterized by other phenotypic traits (reduced motility, reduced antibiotic, haemolytic, and lipolytic activities) (Boemare *et al*., 1997; Cambon *et al*., 2019). Secondary variants display a growth advantage in stationary-phase (GASP) in the insect but are less efficiently transmitted by the nematode than the WT strain (Elhers *et al*., 1990; Cambon *et al*., 2019). The switch to secondary forms in *Xenorhabdus* has been shown to be caused by *lrp* mutations (Cambon et al., 2019), and the *paxAB* gene expression is Lrp-dependent (Engel *et al*., 2017). Indeed, a *lrp* mutant exhibits all the phenotypic traits of a secondary variant, whereas Δ*paxA* mutant share only some of them including BBT adsorption, lecithinase-like activity and abolished PAX production (Cowles *et al*., 2006). Moreover, Lrp is a global activator that affects expression of NRPS genes in *Xenorhabdus* (Jin *et al*., 2024). Our mass spectrometry analysis confirmed that *Xenorhabdus* strains that adsorbed BBT on NBTA medium, were able to produce PAX peptides in culture. Conversely, *X. poinarii* G6, *X. ishibashii* DSM 22670 and secondary variants form red colonies on NBTA medium, exhibit lecithinase-negative phenotypes (Akhurst, 1986) and do not produce PAX peptides in culture (Table S3).

Here, we demonstrated that PAX peptides were related to phenotypes involved in phenotypic variation in *Xenorhabdus* and that the blue phenotype on NBTA medium is a reliable marker for PAX peptide production.

### PAX peptides display multiple roles throughout the life cycle of *Xenorhabdus*

While specialized metabolites produced by soil bacteria have been widely investigated in laboratory conditions for their potential use as antimicrobials in diverse applications, only few studies have addressed their natural roles and little is known about their effective occurrence in natural environments and/or in mutualistic relationships with hosts (Dror *et al*., 2020 ; Shi *et al*., 2018, Gutiérrez-Chávez *et al*., 2021). Pioneer *in vivo* studies on *Xenorhabdus* specialized metabolites demonstrated antimicrobial activity partially attributed to the presence of xenocoumacin in *G. mellonella* larvae infected with *X. nematophila*, with detectable levels persisting for up to 10 days post-infection (Maxwell *et al*., 1994).

Similarly, we monitored by mass spectrometry analysis the presence of *Xenorhabdus* PAX peptides in the insect cadavers in a dynamic way over the course of its natural life cycle. We showed that under simple *in vitro* conditions, PAX peptides were detected from 20 h to 7 days. Under conditions aiming to reproduce the natural life cycle of *X. nematophila* in the insect cadaver (*G. mellonella* infected by *X. nematophila* or infested by *S. carpocapsae*), PAX peptides were detected quite uniformly from 48 h to 7 days post-infection/infestation, but higher variability was observed at 20 h between biological samples. Therefore, PAX peptides are present under natural conditions over a long period of the *Xenorhabdus* life cycle: from the pathogenic phase to the late necrotrophic phase, suggesting that PAX peptides might be involved in various biological functions in *Xenorhabdus*.

PAX peptides may be involved in crucial phenotypes that enable the bacteria to successfully complete the different steps of its infectious cycle, such as bacterial pathogenicity. Contrary to our observations, delayed mortality of *Manduca sexta* larvae had previously been observed when infected with a *ngrA* mutant of *X. szentirmaii* (Ciezki *et al*., 2019). Here, we observed only minor differences between the Δ*paxA* mutant and the WT strain. Previous studies suggested that PAX peptides may protect *Xenorhabdus* against the humoral immune response of insects including AMPs. Because AMPs are cationic peptides, PAX peptides may induce repulsive force due to positive charge at the bacterial cell wall (Vo *et al*., 2021). However, we showed that the MICs of several insect AMPs were similar for the WT and Δ*paxA* or Δ*ngrA* mutants. Therefore, the potential role of PAX peptides during the pathogenic stage could not be inferred from these experiments.

PAX peptides were originally described as antimicrobial peptides (Gualtieri *et al*., 2009), thus they might play a role in inhibiting growth of competing microorganisms introduced in the insect hemocoel upon infection by IJs (Singh *et al*., 2014 ; Ogier *et al*., 2020), or within the cadaver during the necrotrophic stage. However, we showed that nematode microbiota (FAM) (Ogier *et al*., 2020) and insect microbiota displayed low susceptibilities to the three synthetic PAXs tested (PAX1’, PAX2’ and PAX7). PAX peptides therefore do not appear to be required to out-compete microbial competitors.

PAX peptides could be detected up to 7 days post-infection/infestation in our *in vivo* experiment (Figure 1), which is the period when nematodes begin to emerge from the insect cadaver as IJs (Figure S4). Nematode reproductive success assays in *G. mellonella* revealed significant lower progeny using the Δ*paxA* and Δ*ngrA* mutants as compared to the WT strain, demonstrating that PAX peptides are involved in the production of IJs. Previous studies had shown that *ngrA*-dependent compounds were required for the development of the nematode partner in two different nematode genera: *H. bacteriophora* associated with *P. luminescens* and *S. carpocapsae* associated with *X. nematophila* AN6 (Ciche *et a*l., 2001; Singh *et al*., 2015). Our result strongly suggests that PAX peptides are the *ngrA-*dependent metabolites necessary for full IJs production in the *X. nematophila*-*S. carpocapsae* mutualism.

As cyclolipopeptides are often identified as involved in biofilm formation, swimming and swarming motilities (Gutiérrez-Chávez *et al*., 2021), we assessed the involvement of PAX peptides in these phenotypes. Indeed, these features could play a role in successive bacterial colonization of the nematodes in the insect cadaver, prior to IJ emergence. We have highlighted that PAX peptides are positively involved in biofilm formation and negatively involved in swimming motility, but no difference was found in swarming motility. It has been shown that *X. nematophila* can produce biofilms to adhere to the heads of *C. elegans* nematodes, while secondary variants which are unable to produce PAX peptides, or a biofilm-deficient mutant, lacks this ability (Drace *et al*., 2008). Another study carrying out experimental evolution between *Pseudomonas lurida* and *C. elegans* showed that biofilm formation was a bacterial trait associated with a mutualistic lifestyle and motility associated with a free-living lifestyle (Obeng *et al*., 2023). In addition, *P. lurida* produces cyclolipopeptides involved in a mutualistic relationship with the nematode *C. elegans* (Kissoyan *et al*., 2019). We can presume that PAX peptides, which promote biofilm formation, could help the bacterium to associate with its nematode. Other studies revealed that *C. elegans* nematodes could directly sense cyclolipopeptides (serrawettin W2) produced by harmful bacteria (S*erratia marcescens*) with their chemosensory neurons, which had a repulsive effect on the nematodes (Pradel *et al*., 2007). It could be hypothesized that PAX cyclolipopeptides may be sensed by the nematode partner to promote mutualism.

All together, these results revealed that PAX peptides are involved in biofilm formation, swimming motility and the production of nematode IJs.

### The PAX peptide-producing ability is widespread in the genus *Xenorhabdus*

We highlighted that the *paxTABC* cluster was present throughout all the *Xenorhabdus* genus but restricted to that genus only. A distinct genomic environment around *paxTABC* cluster was observed among different *Xenorhabdus* species and a high degree of variability in the percentage of identity within the NRPS gene sequences (*paxABC*) was noticed when blasted to the *X. nematophila* F1 genome. This variability, due to high rate of recombination, is characteristic of NRPS genes, unlike genes encoding proteins via the ribosomal pathway (Fischbach *et al*., 2008 ; Baunach *et al*., 2021 ; Shi *et al*., 2022). We therefore performed the NRPS gene analysis on the modules rather than on the whole genes. Despite NRPS module sequence variability, the 7 different strains representative of the main *Xenorhabdus* species all produced PAX1’ (lysine in position 2), while PAX peptides with an arginine in position 2 (PAX2’ or PAX6) were also detectable in a few strains. In addition to our analysis, the production of PAX peptides has already been confirmed in 8 other strains including *X. nematophila* ATCC 19061, *Xenorhabdus stockiae* DSM 17904, *Xenorhabdus* sp. KJ12.1, *Xenorhabdus* sp. PB62.4, *X. doucetiae* FRM16, *Xenorhabdus khoisanae* SB10, *X. khoisanae* J194 and *X. doucetiae* DSM 17909 (Fuchs *et al*., 2011; Tobias *et al*., 2017; Dreyer *et al*., 2019 ; Booysen *et al*., 2021 ; Vo *et al*., 2021). Nevertheless, it is important to note that not all PAX peptides are detected in all the analyzed strains and the only PAX peptide consistently detected is PAX1’. The ability to produce PAX peptides is therefore both widespread and unique to the *Xenorhabdus* genus.

The only non-PAX-producing species among those tested are *X. ishibashii* and X. *poinarii*. *X. poinarii* has the particularity of being defective in the production of IJs of its nematode partner *Steinernema glaseri*, being avirulent without its nematode and showing strong genomic reduction (Akhurst, 1986; Rosa *et al*., 2002; Ogier *et al*., 2014 ; McMullen *et al*., 2017). Moreover, a large proportion of *S. glaseri* nematodes are naturally aposymbiotic (Akhurst, 1986). It is conceivable that the absence of PAX peptide production by *X. poinarii* may contribute to its inefficient association with its nematode partner. Furthermore, in *X. nematophila* the *paxTABC* genes are clustered close to the *nilABC* genes, whose proteins have been shown to be involved in species-specific colonization of *S. carpocapsae* (Heungens *et al*., 2002; Cowles *et al*., 2008; Bhasin *et al*., 2012; Grossman *et al*., 2022). The *nilABC* cluster is restricted to the *X. nematophila* species, whereas the *paxTABC* genes are present throughout the whole *Xenorhabdus* genus. We can therefore assume that the PAX peptides unique to the *Xenorhabdus* genus are metabolites involved in the mutualistic association between the bacterium and its nematode host.

Finally, PAX peptides unique to the *Xenorhabdus* genus are metabolites involved in many functions such as swimming motility, biofilm formation and the production of infective juvenile nematodes, suggesting their involvement in the interaction between the bacterium and its nematode host.

## Materials and methods

### Bacterial strains, plasmids and growth conditions

The strains and plasmids used in this study are listed in Table S4. Bacteria were routinely grown in Lysogeny Broth (LB, Sigma-Aldrich) medium at 28°C (*Xenorhabdus, S. maltophilia, P. protegens, Ochrobactrum* sp*., Achromobacter* sp*., A. faecalis, Pseudochrobactrum* sp*., Brevundimonas* sp*., E. mundtii* and *D. rugosa*) or 37°C (*E. coli* and *M. luteus*). When required, antibiotics were used at the following final concentrations: gentamicin (Sigma-Aldrich), 20 µg. mL^-1^ and chloramphenicol (Sigma-Aldrich), 20 µg. mL^-1^ for *E. coli* strains and 15 µg. mL^-1^ for *X. nematophila* (or 8 µg. mL^-1^ for allelic exchange).

### Molecular Genetic Techniques

DNA manipulations were performed as previously described (Ausubel *et al*., 1988). Plasmids were introduced into *E. coli* by transformation and transferred to *X. nematophila* by conjugative mating with WM3064 as donor strain (Mouammine *et al*., 2017). All constructs were sequenced by Eurofins Genomics, Germany, GmbH. The primers used in this study (IDT) are described in Table S5.

### Construction of *X. nematophila* Δ*paxA* mutant

The Δ*paxA* mutant was constructed using the same experimental strategy as for the construction of the Δ*ngrA* mutant from the WT strain *X. nematophila* F1 (Lanois *et al*., 2022). The upstream and downstream regions of the *paxA* gene were amplified by PCR with the L-PCR1-*paxA*-SalI and R-PCR1-*paxA*-BamHI primers for the upstream region (621 bp) and the L-PCR2-*paxA*-BamHI and R-PCR2-*paxA*-SpeI primers for the downstream region (601 bp). The two fragments obtained were inserted, together with the 3.8 kb *BamHI* fragment ΩCam cassette from pHP45-ΩCm conferring resistance to chloramphenicol, into pJQ200SK digested with *SalI* and *SpeI* for insertion of the ΩCam cassette between the two PCR fragments. The resulting plasmid, pJQ-*paxA*::ΩCm, was used to transform *E. coli* WM3064 and was introduced into *X. nematophila* F1 by mating. Allelic exchange was performed as previously described (Jubelin *et al*., 2013). ΩCam insertion was confirmed by PCR analysis using primers L-*pax*TA and R-verif-*pax*. The absence of PAX peptide products was assessed by mass spectrometry analysis, as described below. The clone obtained was called Δ*paxA*.

### Phenotypic assays

Phenotypic characteristics of Δ*paxA* and Δ*ngrA* mutant were compared to those of the WT strain by streaking or dropping 5 μL of overnight preculture on the following agar media, as previously described by Boemare *et al*., 1997 : Nutrient Bromothymol Blue Agar (NBTA, Difco Nutrient Agar 1.5% (BD)) supplemented with 25 mg. L^-1^ Bromothymol blue (Merck Millipore) and 40 mg. L^-1^ triphenyl 2.3.5 tetrazolium (TTC, Sigma Aldrich)) ; Difco Nutrient Agar 1.5% supplemented with 0.1 g. L^-1^ CaCl2 and 1% Tween 20 (Polyoxyethylene sorbitan monolaurate), Tween 40 (Polyoxyethylene sorbitan monopalmitate), Tween 60 (Polyoxyethylene sorbitan monostearate), Tween 80 (Polyoxyethylene sorbitan monooleate) or Tween 85 (Polyoxyethylene sorbitan monotrioleate) (Sigma Aldrich) ; Difco Nutrient Agar 1.5% supplemented with 1% egg lecithin (VWR Avantor) ; Trypticase Soja Agar (TSA, BioMerieux) supplemented with 7% sheep blood (Eurobio Scientific) (Vigneux *et al*., 2007) ; Difco Nutrient Agar supplemented with a 6 g. L^-1^ soft agar overlay containing 2% *M. luteus* preculture ; LB 0.35% Agar medium for swimming motility assays (Givaudan *et al*., 1995); and LB supplemented with 0.7% of Eiken Agar (Gerbu) for swarming motility assays (Togashi *et al*., 2000).

### Biofilm assays

Crystal violet biofilm assays were carried out following methods described by Pothula *et al*., 2023, using LB broth instead of LPB medium. Briefly, cultures grown for 48 h without agitation were stained with 1% crystal violet (Sigma Aldrich), then dissolved in 30% acetic acid after washing steps. The absorbance of samples in a 96-well plate was quantified at 550 nm in a TECAN Infinite® 200 plate reader. Each experiment was conducted with four independent clones and repeated four times for each mutant strain. Data were analyzed using GraphPad Prism 9 software and significance was tested using Student’s test.

### MIC determination

MICs were determined in accordance with CLSI guidelines (Clinical and Laboratory Standards Institute, 2015) with following modifications: 5 mL of MHB broth (Bio-Rad) was inoculated with overnight culture in LB broth and grown to a 0.6 < OD_600_ < 0.9. The inoculum for the microplate was prepared as described in the CLSI guide, with dilution in MHB to approximately 10^4^ CFU. mL^-1^. 96-well low-binding plates were incubated for 48 h at 28°C with slow shaking. The peptides used in the analyses were supplied by the following providers: cecropin A (Sigma-Aldrich), cecropin B1 (NeoMPS S.A., Duvic *et al*., 2012), cecropin C (Proteogenix), cecropin P1 (Sigma-Aldrich), drosocin (Eurogentec), colistin (Sigma-Aldrich), polymyxin B (Sigma-Aldrich), NOSO 95-C (odilorhabdin, Nosopharm S.A., Pantel *et al*., 2018). Synthetic PAX1’, PAX2’ and PAX7 were provided by the SynBio3 platform (IBMM, Montpellier) and their structure is shown in Figure S1.

### Pathogenicity assays

Bacterial pathogenicity was assessed by injection of *Xenorhabdus* into *S. littoralis*, as previously described (Givaudan *et al*., 2000). Briefly, bacterial cultures in LB broth (OD_600_ ≈ 0.8) were diluted in the culture medium and 20 μL of the resulting bacterial suspension, containing 1000-3000 CFUs, was injected into the hemolymph of 20 sixth-instar larvae of *S. littoralis*. The number of bacterial cells injected into the larvae was determined by plating on nutrient agar and counting the CFUs. After the bacterial injection, the insect larvae were incubated at 23 °C and mortalities were monitored for up to 50 h. The pathogenicity of bacterial isolates was determined by measuring the time required for 50% of the insect larvae to be killed (LT_50_). The results of 4 independent experiments were combined and analyzed with the software Statistical Package for the Social Sciences V18.0 (SPSS). Significant differences between two datasets were assessed with non-parametric Wilcoxon-Mann-Whitney tests, with a 95% confidence interval.

### Nematode colonization assays

Bacterium–nematode complexes were produced, using an *in vivo* technique (Sicard *et al*., 2004; Jubelin *et al*., 2011) in which *G. mellonella* (wax moth) larvae were infected with *X. nematophila* F1 strain and aposymbiotic IJs of *S. carpocapsae* (SK27, Plougastel) nematodes. Aposymbiotic nematodes were obtained previously (Huot *et al*., 2020). Approximatively 100 aposymbiotic IJs per larvae were first placed on a piece of filter paper in individual Eppendorf tubes and incubated with 20 larvae of *G. mellonella* at 23°C. After 24 h of infestation, insect larvae were injected with 20 µl (∼3000 bacteria) of WT, Δ*paxA* or Δ*ngrA* strains that had been grown until OD_600_ ≈ 0,8. After insect death (24 to 48 h post infestation), cadavers were transferred to White traps (White, 1927). Fourteen and thirty days after infestation, the IJs emerging from the cadavers were collected in Ringer’s solution (Aguettant). Once again, these IJs were then used to infest new *G. mellonella larvae*. Nematodes were collected and washed with water through a 20 µM filter and stored in Ringer at 9°C in tissue culture flasks. Reproductive success (number of IJ nematodes emerging from an insect larva) was determined by cumulating the number of nematodes collected and counted at 15- and 30-days post-infestation. Direct counts of average CFU/nematode were determined via a grinding assay wherein collected nematodes were equalized by density, serially diluted, and plated on NBTA plates. Each experiment was repeated three times. Data were analyzed using GraphPad Prism 9 software and significance was tested using non-parametric Wilcoxon-Mann-Whitney tests.

### Mass spectrometry analysis

#### PAX peptide extraction from *Xenorhabdus* cultures

PAX extraction methods were adapted from those previously described by Gualtieri *et al*., 2009, Fuchs *et al*., 2011 and Vo *et al*., 2021. Tubes of 5 mL LB were inoculated with 200 µL preculture of overnight *Xenorhabdus* strains and grown for 48h at 28°C under agitation. Cultures were centrifuged at 6000 x *g*, 10 min and the supernatant fraction were discarded. The cell pellet fraction was resuspended in 1 mL LB and 45 µg of DP18 (R-3-Hydroxytetradecanoic acid-Gly-Orn-[DLys-Lys-DLys-DLys-Lys], m/z=1038.92 Da), a synthetic PAX peptide derivative, was added as an extraction control. Samples were sonicated for 2 min, then centrifuged at 16000 x *g*, 5 min. The supernatant corresponding to the cytosolic fraction (S1) was conserved. The new pellet containing membrane debris was resuspended in 500 μl methanol/H_2_O (vol/vol) and acidified with 1% formic acid. Samples were incubated for 5 min in the ultrasonic bath. The insoluble fractions were then separated by centrifugation (16000 x *g*, 5 min). The resulting supernatant S2 was pooled with supernatant S1 and then filtered with 0.45 µM filters (Filtropur, Sarstedt). Samples were suspended at a 1:1 ratio in 0.1 M NaCl, 0.02 M Tris, pH 9 (vol/vol), then loaded onto Sep-Pak CarboxyMethyl Short Cartridge (Accell Plus CM, Waters) prepared according to the manufacturer’s instructions. Cartridges were eluted with 0.5 M NaCl, 0.02 M Tris, pH 9, 0.1% trifluoroacetic acid (TFA). The resulting peptide extract was then loaded onto Sep-Pak C18 Short cartridge (Sep-Pak Plus C18, Waters) prepared according to the manufacturer’s instructions. Peptides were eluted with 100% acetonitrile (ACN) and freeze-dried.

#### Preparation of *in vivo* samples

*G. mellonella* larvae were injected with 5.10^3^ to 7.10^3^ CFUs of *X. nematophila* F1 (as described methods in Pathogenicity assays) or infested with 100 *S. carpocapsae* SK27 IJs nematodes (as described methods in Nematode colonization assays) and incubated at 23°C. Larvae were weighed and frozen at -20°C at each time point post-injection/infestation: 20 h, 48 h, 72 h, 5 days and 7 days. Larvae surfaces were disinfected with ethanol and cuticles were removed. The insides of 10 larvae per condition were pooled and added to 2 mL 0.05 M Tris, pH 7 before grinding using FastPrep®-24 5G (MP Biomedicals) with sterile glass beads (6.5 m/s, 30 s, x3). Samples were centrifuged at 6000 x *g*, 30 min and cell pellets were resuspended in 1.5 mL 0.05 M Tris, pH 7. PAX peptide extraction was then performed as described above, starting from the step of DP18 addition to the samples. Experiments were performed in triplicate.

### MALDI-TOF-MS Analysis

Extracted peptides were analyzed with a RapifleX MALDI TOF/TOF (Bruker) equipped with a Pulse Smart Beam 3D laser at 335 nm. The following instrument parameters were used: frequency, 10 000 Hz; delayed extraction time, 160 ns; source, positive mode; reflectron mode. Samples were resuspended in ACN and mixed at a 1:1 ratio with 1 μL of saturated α-cyano-4-hydroxycinnamic acid (CHCA). 1 μL of the mix was filed onto a stainless-steel target and air-dried. A targeted analysis of the spectra was carried out on compounds with m/z ratios between 1000 and 1200 and corresponding monoisotopic mass lists were generated using Flex Analysis 3.4 (Bruker).

### NRPS genomic analysis

The *paxTABC* and surrounding gene sequences of 40 strains among 23 *Xenorhabdus* species were extracted from the MicroScope MaGe database (https://mage.genoscope.cns.fr/microscope/home/index.php). Module sequences of NRPS genes (*paxA* : m1 ; *paxB* : m2, m3, m4 ; *paxC* : m5, m6, m7) were extracted using the antiSMASH tool implemented in MicroScope MaGe database. Strains with genomes of poor assembly quality (>350 contigs) were removed from the analysis. A blastn analysis using the NCBI BLAST+ tool implemented on the Galaxy v.2.10.0+galaxy1 platform (Galaxy Pasteur, https://galaxy.pasteur.fr/) was performed using the sequence of *X. nematophila* F1 as query with a cutoff value of 0.1 and a minimum coverage percentage >30%. Percentage identities have been used to construct Figure 6 using ggplot2 package on RStudio v2024.09.1+394. Details of the data used for these analyses can be found in Tables S6 and S7.

## Acknowledgments

We acknowledge the Insectarium Support Platform, DGIMI, for providing *G. mellonella* and *S. littoralis* larvae. We thank Elodie Carretero and Pascal Verdie from the SynBio3 platform (IBMM, UMR CNRS 5247, Montpellier) supported by GIS IBISA, for synthesis of PAX peptides. This work was supported by the SPE department of INRAe, through the AntibioNEP (IB2023-AAPSPE) project, and by the RIVOC Key Challenge (PFI 201555PI), funded by the Occitanie Region and led by the University of Montpellier (UM). Noémie Claveyroles was supported by a doctoral fellowship from the Doctoral School Gaia, UM. Imane El Fannassi was the recipient of a Masters scholarship from LabUM chimie, UM.

